## Supplementary Material for "Markov State Modelling Reveals Heterogeneous Drug-Inhibition Mechanism of Calmodulin"

*Supplementary Information*

Annie M. Westerlund[1†], Akshay Sridhar[1†], Leo Dahl[1], Alma Andersson[1,2], Anna-Yaroslava Bodnar[1], Lucie Delemotte[1*]

[1] Department of Applied Physics, Science for Life Laboratory, KTH Royal Institute of Technology, 17121, Solna, Sweden

[2] Division of Gene Technology, Science for Life Laboratory, KTH Royal Institute of Technology, 17121, Solna, Sweden

[†] Contributed equally to the work

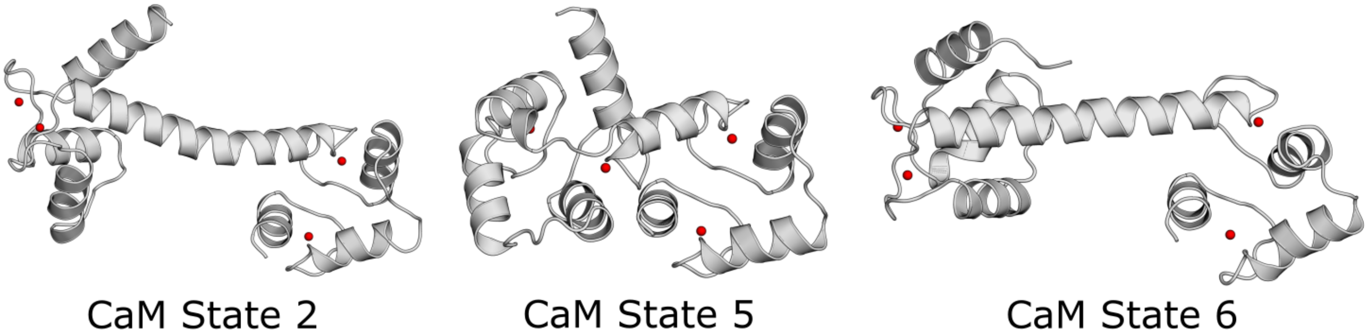


**Figure 1** Calmodulin states 2, 5 and 6 obtained from Molecular Dynamics simulations of the 3CLN structure that were used as initial configurations for the REST simulations. The bound $Ca^{2+}$ ions included in the simulations are shown as red spheres.

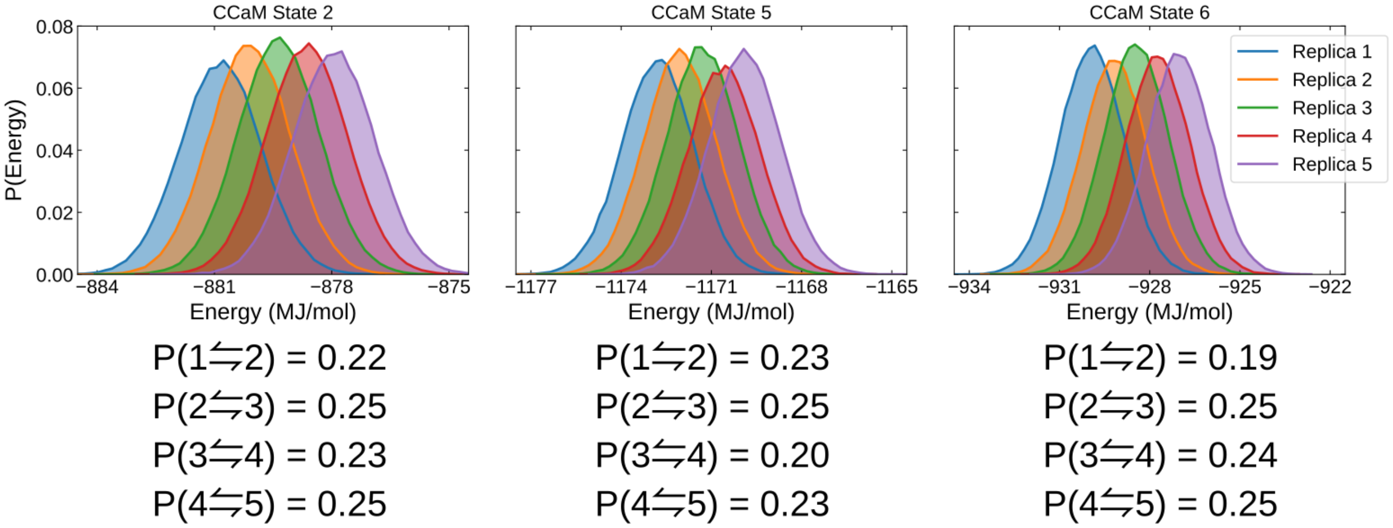


**Figure 2** Efficiency of the Replica-Exchange with Solute Tempering (REST) simulations initiated from the three CaM states assessed by the energy overlap and mean exchange acceptance probability between adjacent replicas.

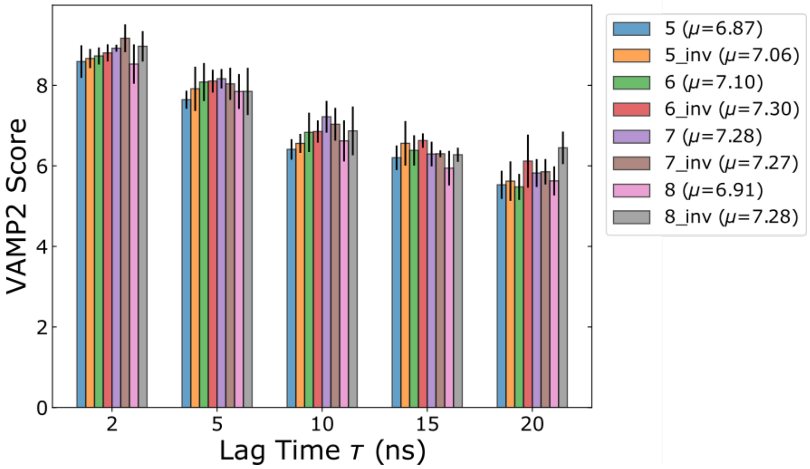


**Figure 3** VAMP2 scores of the 10 slowest processes for different feature transformations calculated at a variety of lag times $\tau$. The mean values from five-fold cross-validation are plotted as bars and error bars represent the standard deviations. The mean value across the lag times calculated for each feature transformation is mentioned within the legend.

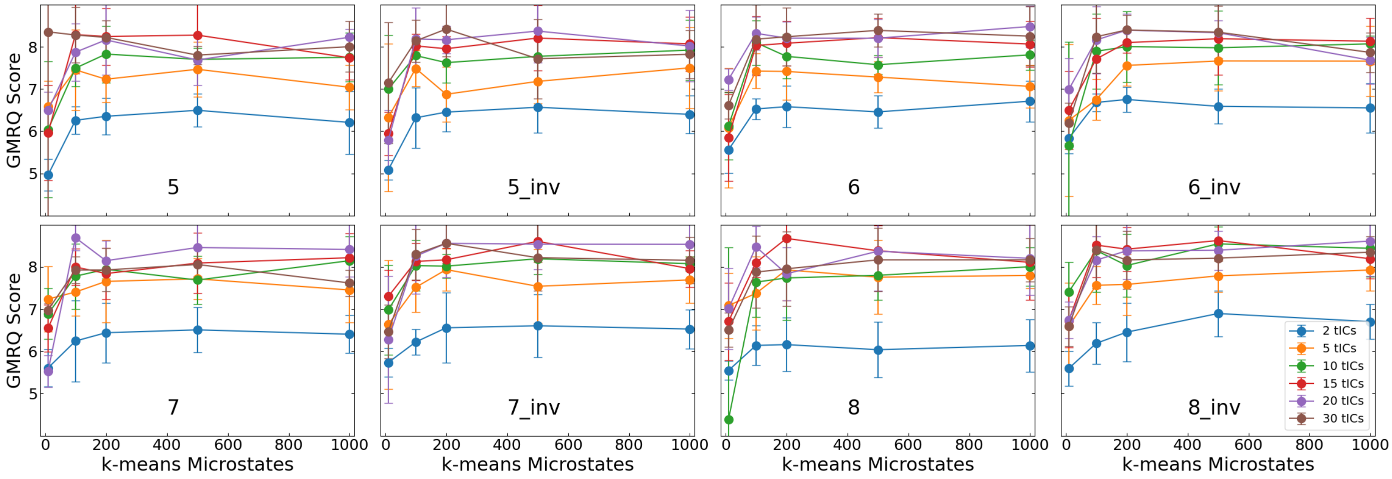


**Figure 4** The optimization of MSM hyperparameters through the calculation of GMRQ scores for different feature transformations at varying numbers of microstates and processes. The mean values from five-fold cross-validation are plotted as dots and standard deviations are plotted as error bars.

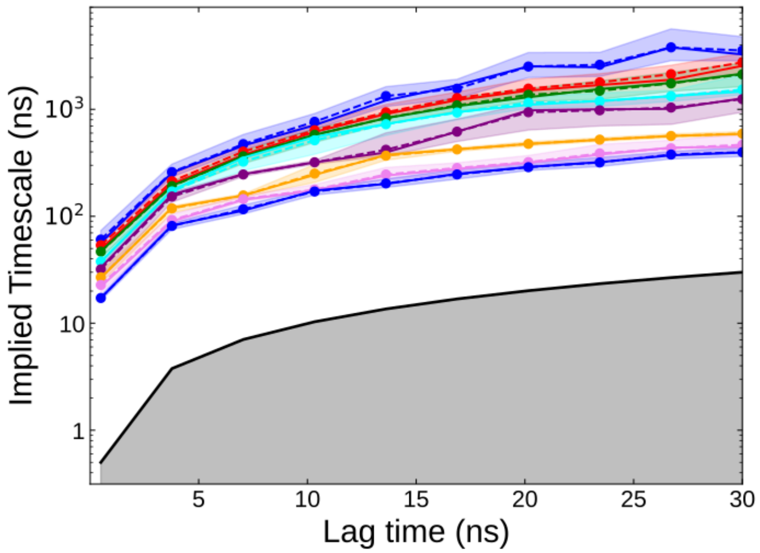


**Figure 5** Top 8 eigenvalues of the transition probability matrix calculated at varying lag-times to identify a memoryless Markovian time. The 95% confidence intervals of the eigenvalues are shown as shaded regions. The black solid curve delimits a shaded region where the implied timescales are shorter than the lagtime.

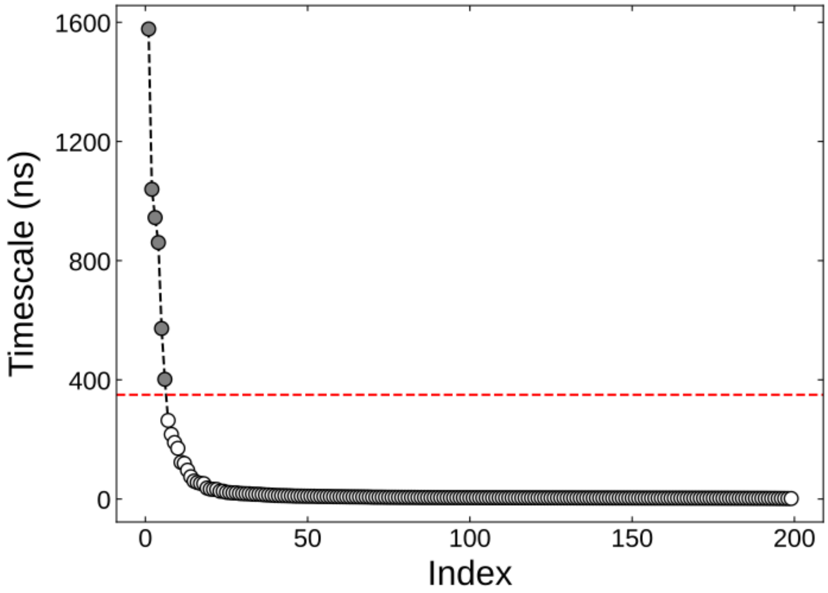


**Figure 6** Spectral analysis of the eigenvalues to identify the number of clusters. The cutoff between the sixth and seventh relaxation timescales selected in this work is illustrated as red dotted line.

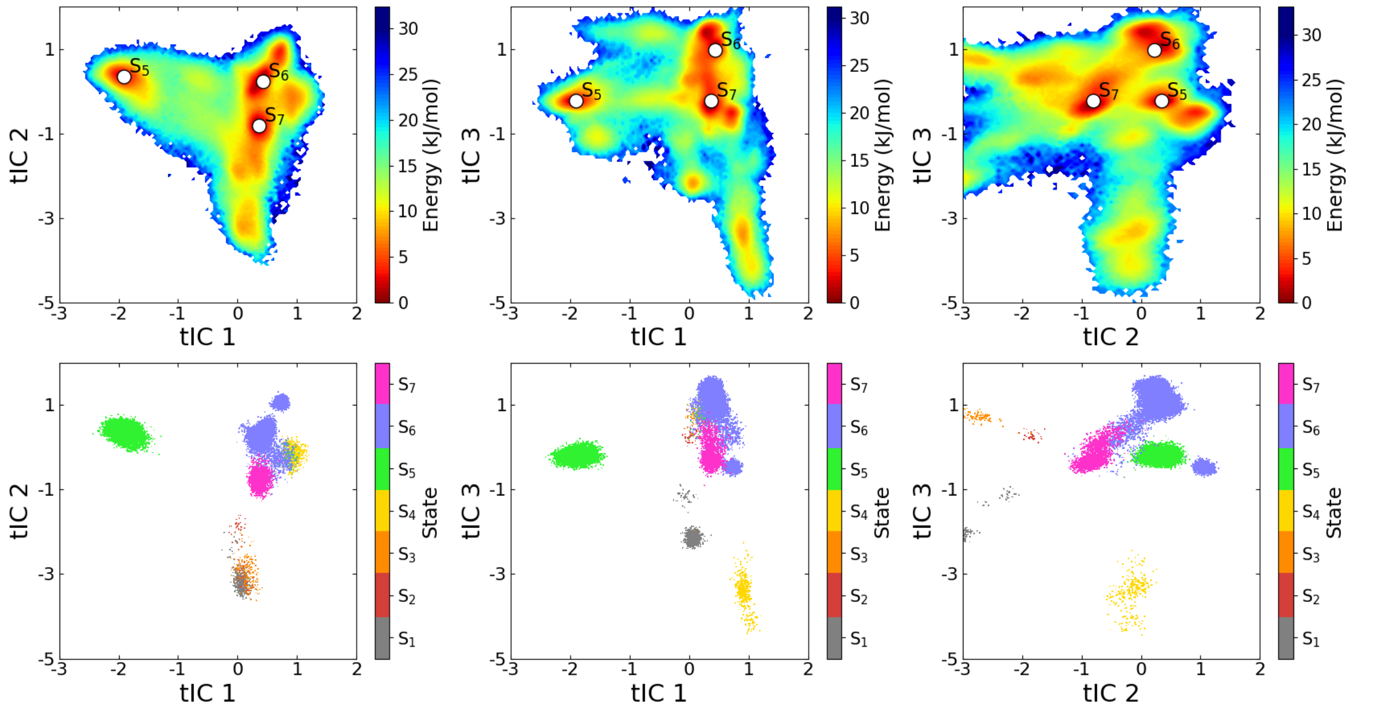


**Figure 7 Top:** Projection of the Markov state model free-energy surface along the three slowest time-lagged independent components (tICs). **Bottom:** $S_1$-$S_7$ macrostate assignments along the three tICs obtained from PCCA++ clustering. Each trajectory frame is represented as a dot within the scatter plot.

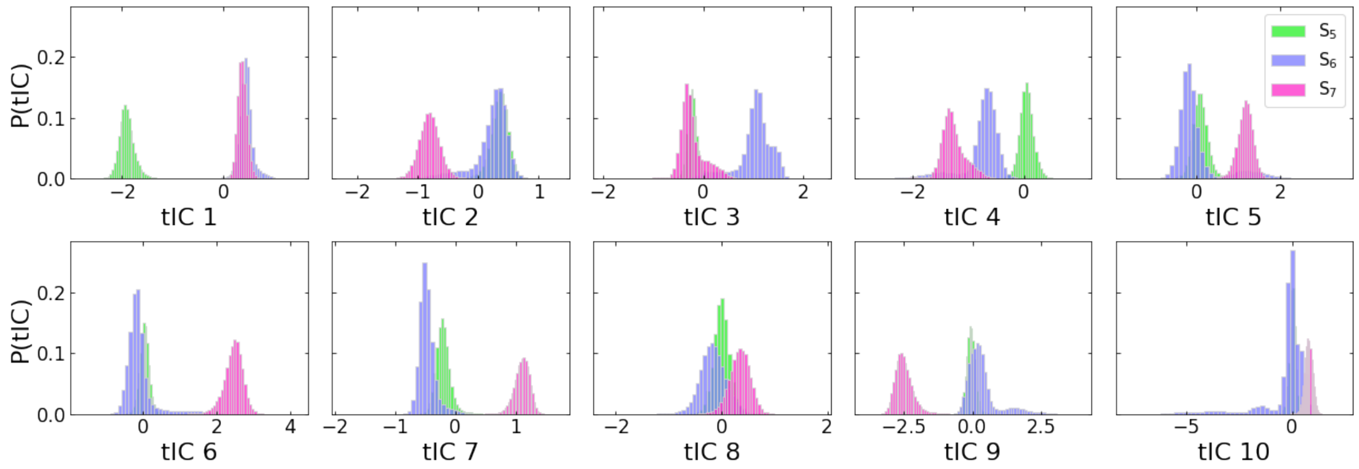


**Figure 8** Separation of the $S_5$, $S_6$ and $S_7$ macrostates along the 10 slowest time-lagged independent components (tICs) used for Markov state model construction.

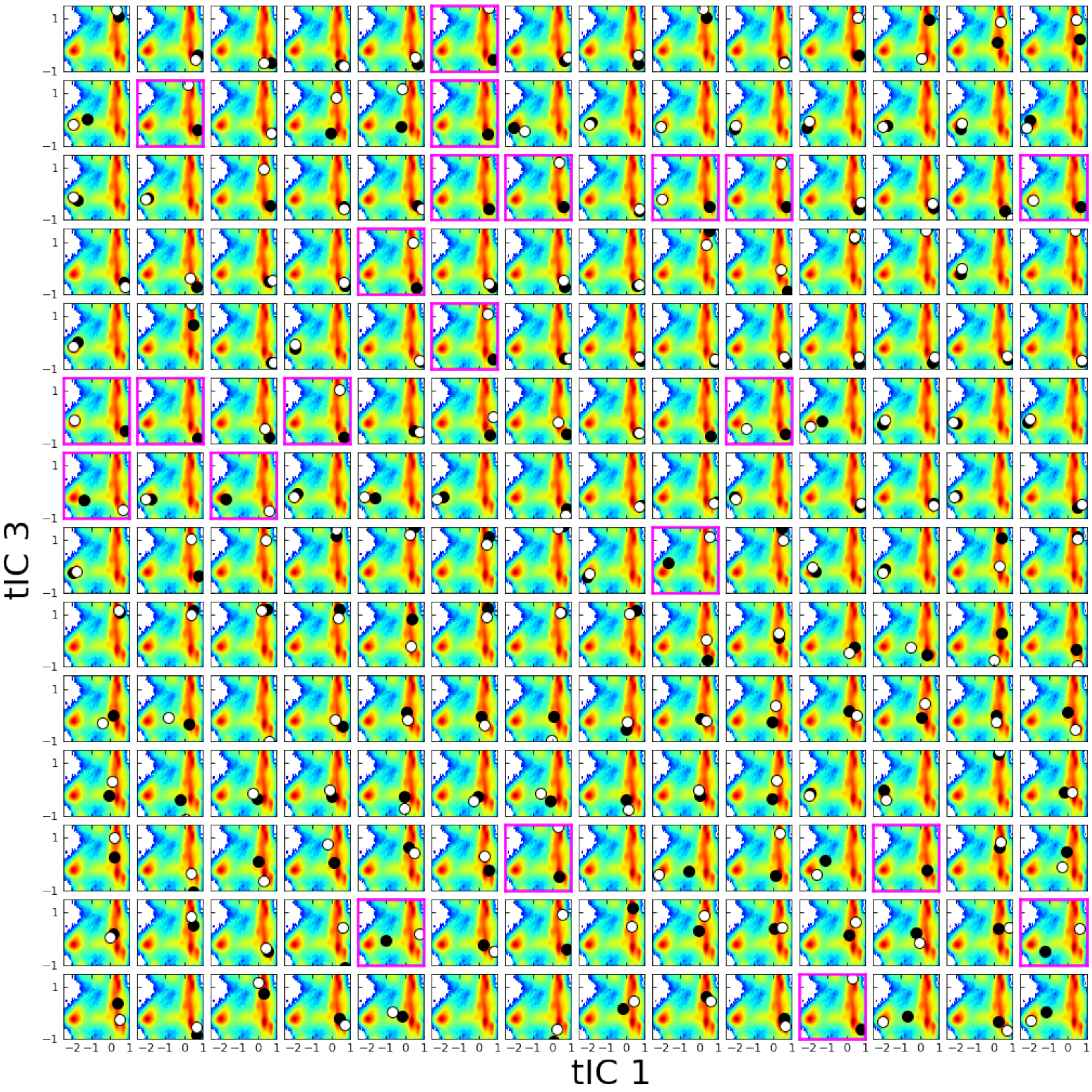


**Figure 9** Transitions between the macrostate basins analyzed by projecting the initial and final configurations of individual trajectories onto the free-energy surface. Individual trajectories are represented as subplots with the initial and final configuration shown as black and white dots respectively. Trajectories with transitions between the basins are highlighted with a magenta outline.

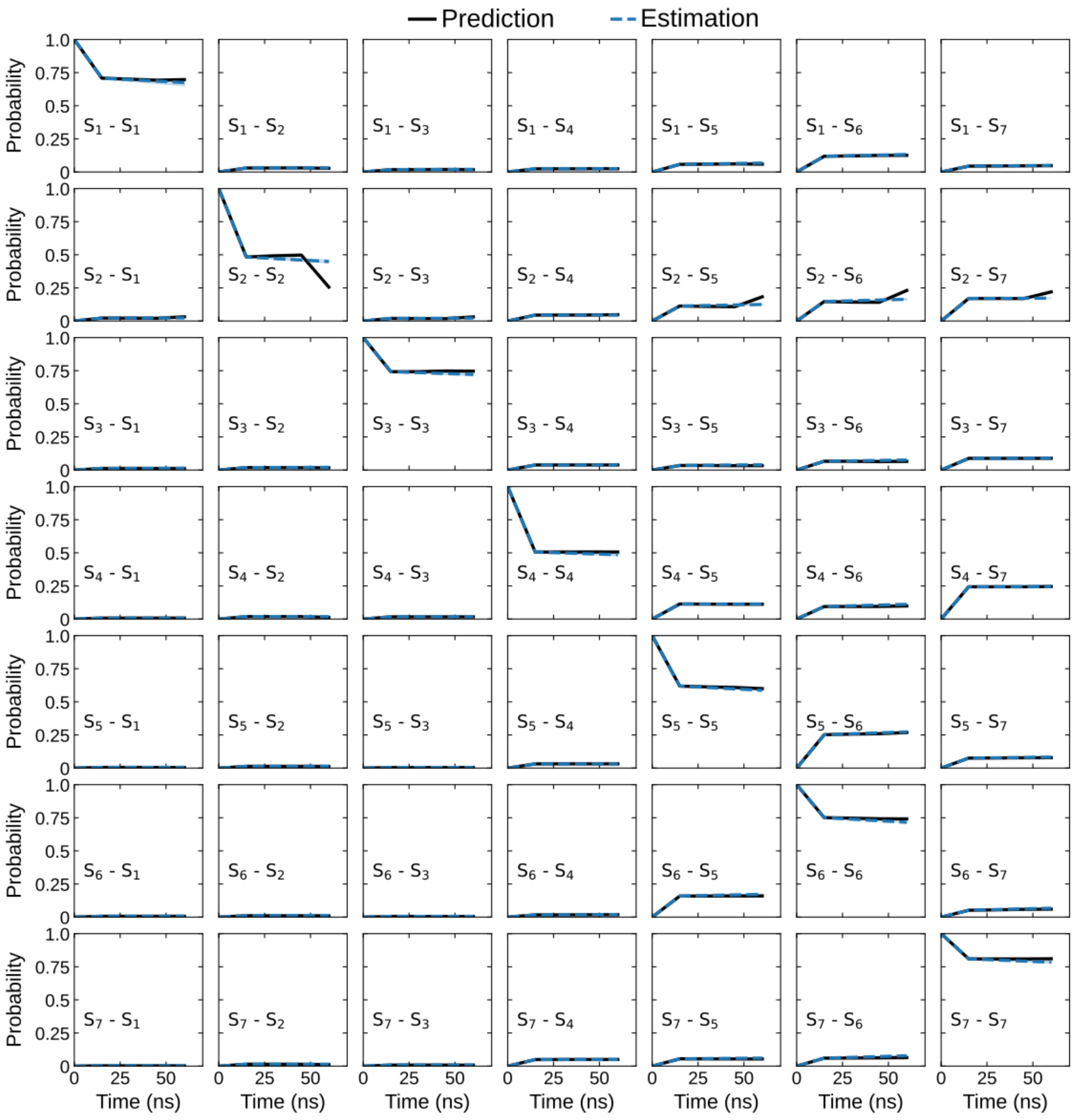


**Figure 10** Chapman-Kolmogorov test validating the Markov state model by comparing the probabilities of transiting between the macrostates (blue line) and the calculated probabilities from the constructed model (black line).

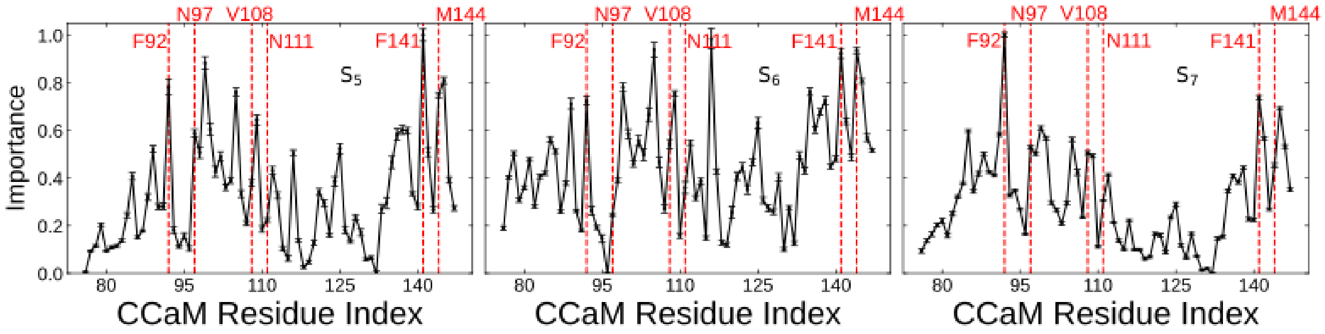


**Figure 11** The per-residue importance in discerning the $S_5$, $S_6$ and $S_7$ macrostates calculated using the supervised KL Divergence method. Plots represent the mean values calculated from five-fold cross-validation and the standard deviations are plotted as error bars. Physiologically important residues and those with high importance values are illustrated using red dotted lines.

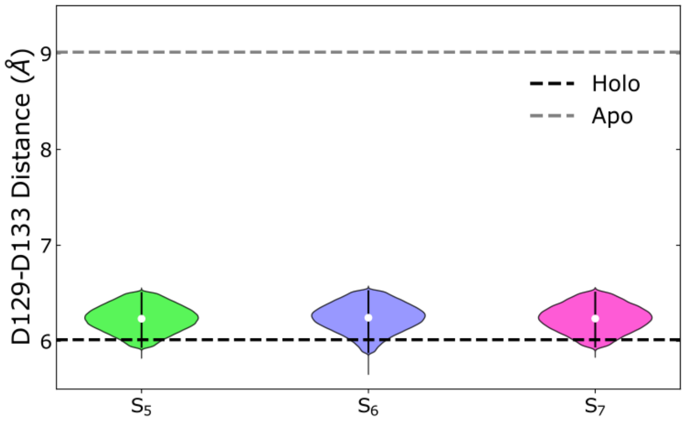


**Figure 12** Distance between center-of-mass of the D129 and D133 acidic residues making up the second $Ca^{2+}$ binding site within the $S_5$, $S_6$ and $S_7$ macrostates. Each violinplot spans the 5th and 95th percentile of distances and is weighted by the Markov state model probabilities. The median value for each macrostate is represented as a white dot. The inter-residue distances calculated from *apo-* (PDB: 1CFD) and *holo-* (PDB: 1CLL) calmodulin structures are shown as grey and black dotted lines respectively.

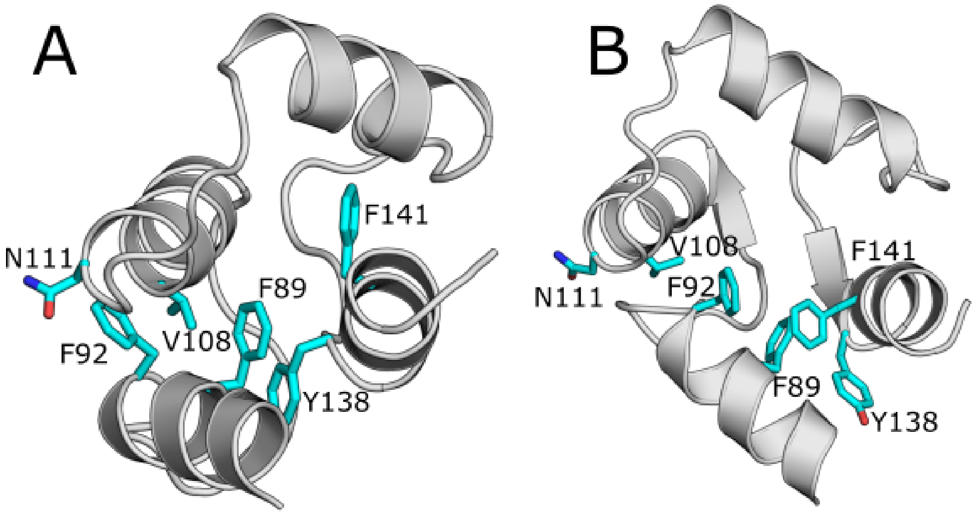


**Figure 13** Transition in the stacking of aromatic residues between the **(A)** *apo-* (PDB: 1CFD) and **(B)** *holo-* (PDB: 1CLL) states of calmodulin induced by the binding of $Ca^{2+}$ ions.

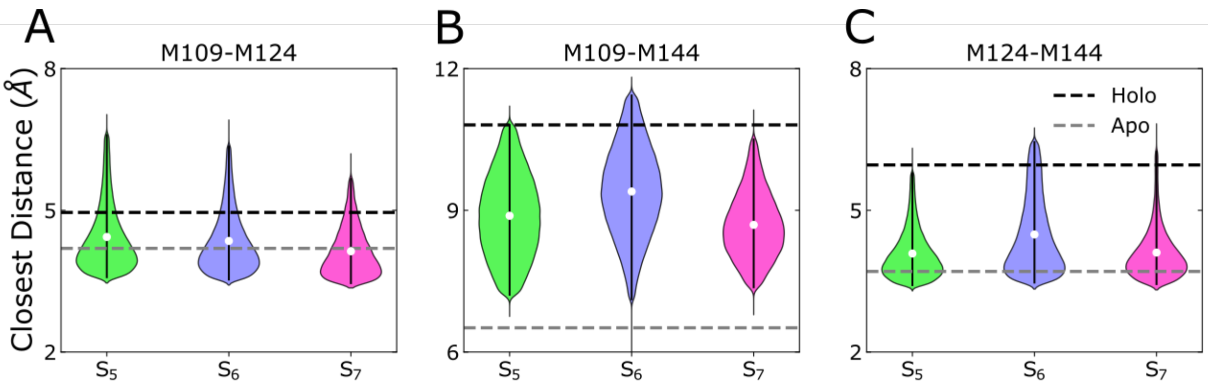


**Figure 14** Distances between the sidechains of the M109, M124 and M144 Methionine residues making up the hydrophobic binding pocket within the $S_5$, $S_6$ and $S_7$ macrostates. Each violinplot spans the 5th and 95th percentile of distances and is weighted by the Markov state model probabilities. The median value for each macrostate is represented as a white dot. The inter-residue distances calculated from *apo*- (PDB: 1CFD) and *holo*- (PDB: 1CLL) calmodulin structures are shown as grey and black dotted lines respectively.
